# Byproducts of inflammatory radical metabolism provide transient nutrient niches for microbes in the inflamed gut

**DOI:** 10.1101/2023.12.08.570695

**Authors:** Luisella Spiga, Maria G. Winter, Matthew K. Muramatsu, Vivian K. Rojas, Rachael B. Chanin, Wenhan Zhu, Elizabeth R. Hughes, Savannah J. Taylor, Franziska Faber, Steffen Porwollik, Tatiane F. Carvalho, Tian Qin, Renato L. Santos, Helene Andrews-Polymenis, Michael McClelland, Sebastian E. Winter

## Abstract

Louis Pasteur’s experiments on tartaric acid laid the foundation for our understanding of molecular chirality, but major questions remain. By comparing the optical activity of naturally-occurring tartaric acid with chemically-synthesized paratartaric acid, Pasteur realized that naturally-occurring tartaric acid contained only L-tartaric acid while paratartaric acid consisted of a racemic mixture of D- and L-tartaric acid. Curiously, D-tartaric acid has no known natural source, yet several gut bacteria specifically degrade D-tartaric acid. Here, we investigated the oxidation of monosaccharides by inflammatory reactive oxygen and nitrogen species. We found that this reaction yields an array of alpha hydroxy carboxylic acids, including tartaric acid isomers. Utilization of inflammation-derived D- and L-tartaric acid enhanced colonization by *Salmonella* Typhimurium and *E. coli* in murine models of gut inflammation. Our findings suggest that byproducts of inflammatory radical metabolism, such as tartrate and other alpha hydroxy carboxylic acids, create transient nutrient niches for enteric pathogens and other potentially harmful bacteria. Furthermore, this work illustrates that inflammatory radicals generate a zoo of molecules, some of which may erroneously presumed to be xenobiotics.

## INTRODUCTION

Host-associated microbial communities fulfill critical functions beneficial to their host, such as immune education, protection from enteric pathogens, and regulation of energy metabolism ^1,2^. Under homeostatic conditions, the population size of each microbial species is controlled by nutrient availability, the capacity of each microbe to access specific nutrients, and nutritional competition (nutrient niche) ^3^. With most monosaccharides and amino acids being absorbed in the small intestine, complex poorly absorbed molecules such as polysaccharides are major carbon and energy sources for colonic microbes. The genetic capacity to degrade glycans correlates with the abundance of any given microbe in the colon ^4,5^. Inclusion of a single molecule in the diet can create a new nutrient niche filled by specific microbes. For example, administration of a marine porphyran in the diet creates a private nutrient niche and allows for stable colonization with a porphyran-degrading *Bacteroides* strain ^6^. The food additive trehalose supports growth of epidemic *C. difficile* and increases virulence in a mouse model ^7^. While the composition of the gut microbiota is altered in a number of diseases, the underlying changes in nutrient niches that support the growth of potentially harmful bacteria are not completely understood.

Many nutrients consumed by gut bacteria in the large intestine are derived from plant material, such as tartaric and other, related α-hydroxycarboxylic acids. L-tartaric acid is generated as part of ascorbate metabolism via the Smirnoff-Wheeler pathway in grapevines (*Vitis spp.*) ^8,9^. In addition, L-tartrate forms the backbone for chicoric acid produced by endive (*Cichorium spp.*) and coneflowers (*Echinacea spp.*) ^10^. Humans do not metabolize tartrate and most dietary L-tartrate passes into the large intestine ^11^, where it could be utilized by the microbiota. *E. coli* and *Clostridia* spp, *C. rodentium, Klebsiella pneumoniae,* and *Salmonella enterica* serovar Typhimurium (*S*. Tm), are capable of degrading and L-tartaric acid ^12-17^. Curiously, some gut bacteria only metabolize D-tartrate (or both enantiomers) ^12-17^ (**Fig. S1A).** Yet there are no known natural sources of D-tartrate, suggesting that natural sources of D-tartrate may indeed exist but have eluded discovery.

Reactive oxygen species (ROS) and nitrogen species(RNS) play an important role in shaping host-microbe interactions, by killing bacterial cells and by creating nutrient niches. Cell-type specific NADPH oxidases, superoxide dismutases, and myeloperoxidase generate bactericidal ROS, while RNS are generated during inflammation primarily through inducible nitric oxide synthase (iNOS)^18,19^. Chronic granulomatous disease patients, individuals who are unable to produce necessary amounts of inflammatory ROS, are highly susceptible to opportunistic bacterial and fungal infections, illustrating the importance of this host defense mechanism^20^. In addition to their roles in fighting infections, ROS are generated byproducts of normal mitochondrial respiration and they can also function as cell signaling molecules ^21^. ROS and RNS readily react with a variety of biomolecules and cellular components. Damage to microbial and host cells occurs through specific mechanisms, such as DNA damage, lipid peroxidation, oxidation of dGTP and dCTP pools, as well as protein modification and fragmentation ^22,23^. During intestinal inflammation, ROS and RNS leak into the gut lumen and give rise to electron acceptors for microbial respiration ^24-26^. Apart from these specific examples, we have an incomplete understanding of how inflammatory ROS and RNS react with biomolecules, influence niche creation in the gut, and impact microbial metabolism. In the current study, our investigations on bacterial tartrate metabolism in the mammalian gut reveal an unexpected molecular link between ROS and RNS metabolism, and the generation of α-hydroxycarboxylic acids, such as tartrate, and the growth of Enterobacteriaceae family members.

## RESULTS

### *S.* Tm degrades D- and L-tartrate through two independent, stereospecific pathways

We reasoned that *S*. Tm’s ability to degrade both D- and L-tartrate would allow us to investigate bacterial tartrate metabolism in the mammalian intestinal tract. During anaerobic growth, *S*. Tm ferments both D- and L-tartrate, but not *meso*-tartrate (**Fig. 1A; Fig. S1A and B**)(data not shown). The *S*. Tm genome encodes a multi-subunit dehydratase (*ttdBA*) with homology to the tartrate dehydratase in *E. coli*, as well as a putative transporter (renamed *ttdU*) and putative transcriptional regulators (**Fig. 1B; Fig. S2A**) ^27,28^. To investigate the roles of these genes in L-tartrate utilization, we constructed mutants lacking *ttdBAU* genes. We then cultured an equal mixture of the *S*. Tm wild-type strain and the isogenic *ttdBAU* mutant under anaerobic conditions and determined the ratio of wild-type bacteria to mutant bacteria to generate a competitive index. The wild-type strain outcompeted the *ttdBAU* mutant in the presence of L-tartrate, but not D-tartrate or other small organic acids (**Fig. 1C; Fig. S2B**). Similarly, a mutant lacking the tartrate dehydratase (*ttdBA* mutant) or the transporter (*ttdU*) displayed a competitive fitness defect in media supplemented with L-tartrate, but not D-tartrate (**Fig. 1D and E; Fig. S2C**). In *E. coli*, DcuB is required for the transport of D-tartrate ^29^, while in *Salmonella*, *dcuB* was dispensable for D-tartrate utilization (**Fig. S2D**). These findings suggest that L-tartrate utilization requires the *ttdBAU* genes and that an unknown pathway is responsible for D-tartrate utilization in *S*. Tm.

**Figure 1:**
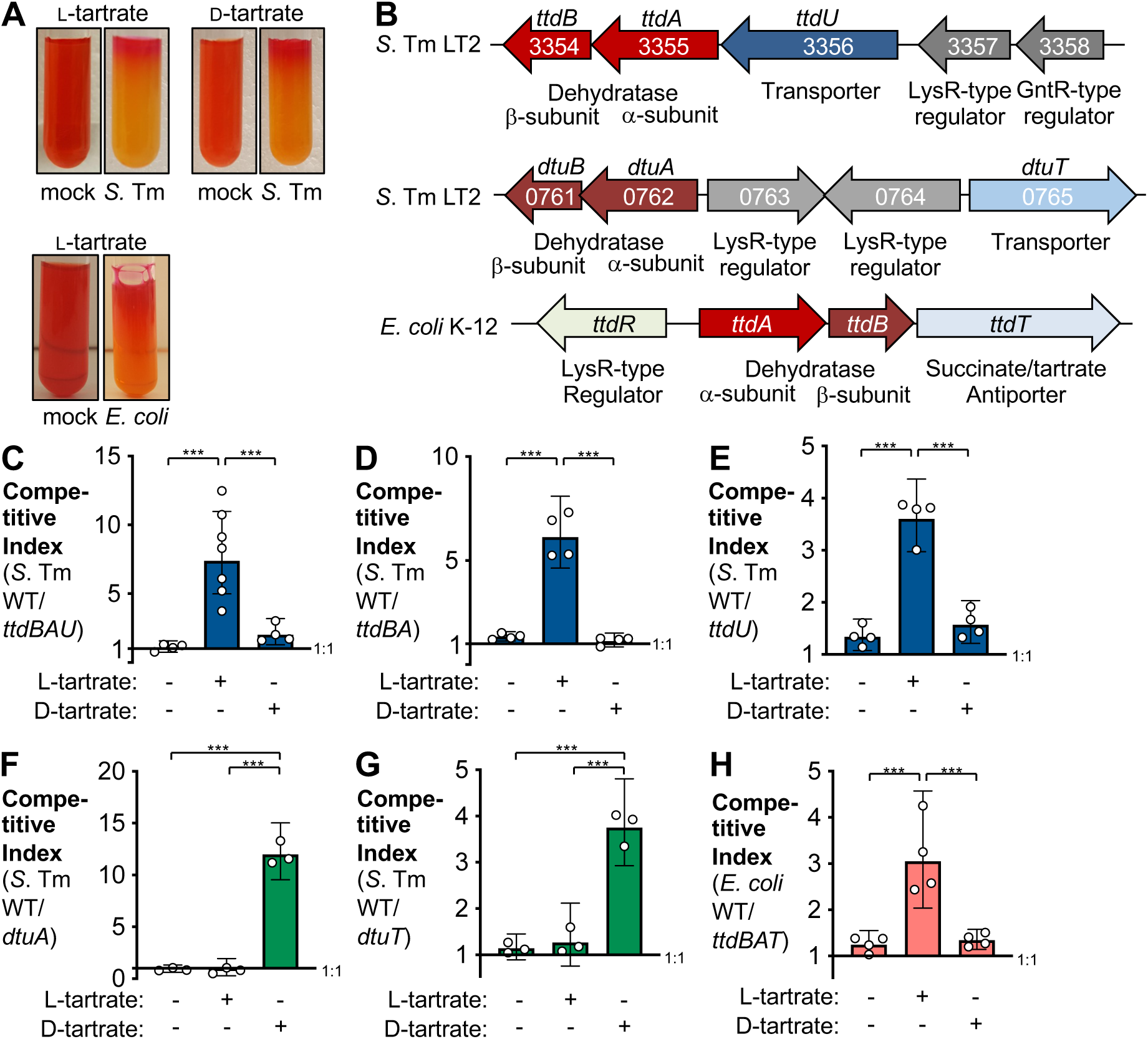
Utilization of tartaric acid by *S. Tm* and *E. coli*. (**A**) Jordan’s agar base, supplemented with either L-tartrate or D-tartrate, was inoculated either with the *S.* Tm wild-type strain or the *E. coli* wild-type strain and incubated at 37 °C overnight. (**B**) Schematic representation of the L-tartrate and the D-tartrate operons in *S.* Tm, and the Ltartrate operon in *E. coli*. **(C-H)** Competitive anaerobic growth in modified M9 medium in the absence or presence of Ltartrate (20 mM) and D-tartrate (20 mM) of the *S.* Tm wild-type strain and a *ttdBAU* mutant (**C**) or a *ttdBA* mutant (**D**) or a *ttdU* mutant (**E**) or a *dtuA* mutant (**F**) or a *dtuT* mutant (**G**), and competitive index of the *E. coli* NRG857c AIEC wild-type strain and an isogenic *ttdBAT* mutant (**H**). The competitive index was determined after 16 hr of anaerobic growth. Each white dot represents one biological replicate. Data are shown as geometric mean +/- geometric standard deviation; * * * *P* < 0.001.

Analysis of predicted coding sequences in the *S*. Tm genome revealed a locus (STM0761-5) with weak similarity to the *ttd* gene cluster (**Fig. 1B**). This candidate locus is comprised of a predicted dimeric dehydratase (STM0761-2; renamed *dtuAB*), two putative transcriptional regulators (STM0763-4), and a putative cation transporter (STM0765; renamed *dtuT*). Mutants lacking *dtuA* or *dtuT* were unable to utilize D-tartrate, and displayed no fitness defect in L-tartrate utilization (**Fig. 1F and G; Fig, S2E**), suggesting that the products of the *dtu* locus mediate D-tartrate utilization.

### Contribution of D- and L-tartrate utilization to fitness of *S.* Tm in the murine gut

Next, we sought to determine whether *S*. Tm utilizes D- and L-tartrate during infection. Infection of immunocompetent individuals with non-typhoidal *Salmonella* strains results in acute gastroenteritis. CBA mice infected with *S*. Tm develop cecal and colonic inflammation, with a significant infiltration by neutrophils ^30,31^. We intragastrically inoculated groups of CBA mice with an equal mixture of the *S*. Tm wild-type and either an isogenic *ttdBAU* mutant or an isogenic *dtuT* mutant. Four days after infection, we determined the fitness (competitive index) of each strain by plating on selective agar (**Fig. 2A-C**). We recovered the wild-type strain in significantly higher numbers than the *ttdBAU* mutant (**Fig. 2A**). Curiously, the *S*. Tm wild-type strain also outcompeted the *dtuT* mutant (**Fig. 2C**), suggesting that both D- and L-tartrate utilization confer a fitness advantage in the murine intestinal tract. When groups of mice were intragastrically inoculated with the wild-type strain, a *ttdBAU*, or a *dtuT* mutant, the D- and L-tartrate utilization-deficient colonized the colon lumen to a lesser extent than the wild-type strain (**Fig. 2D**), indicating that tartrate utilization not only enhances competitive fitness but also contributes to gut colonization. Genetic complementation restored the fitness of the *ttdBAU* and *dtuT* mutants (**Fig. 2A and C**). Collectively, the results demonstrate that D- and L-tartrate utilization both contribute to intestinal colonization by *S*. Tm and suggest that both D- and L-tartrate are present in the murine gut lumen during *S*. Tm infection.

**Figure 2:**
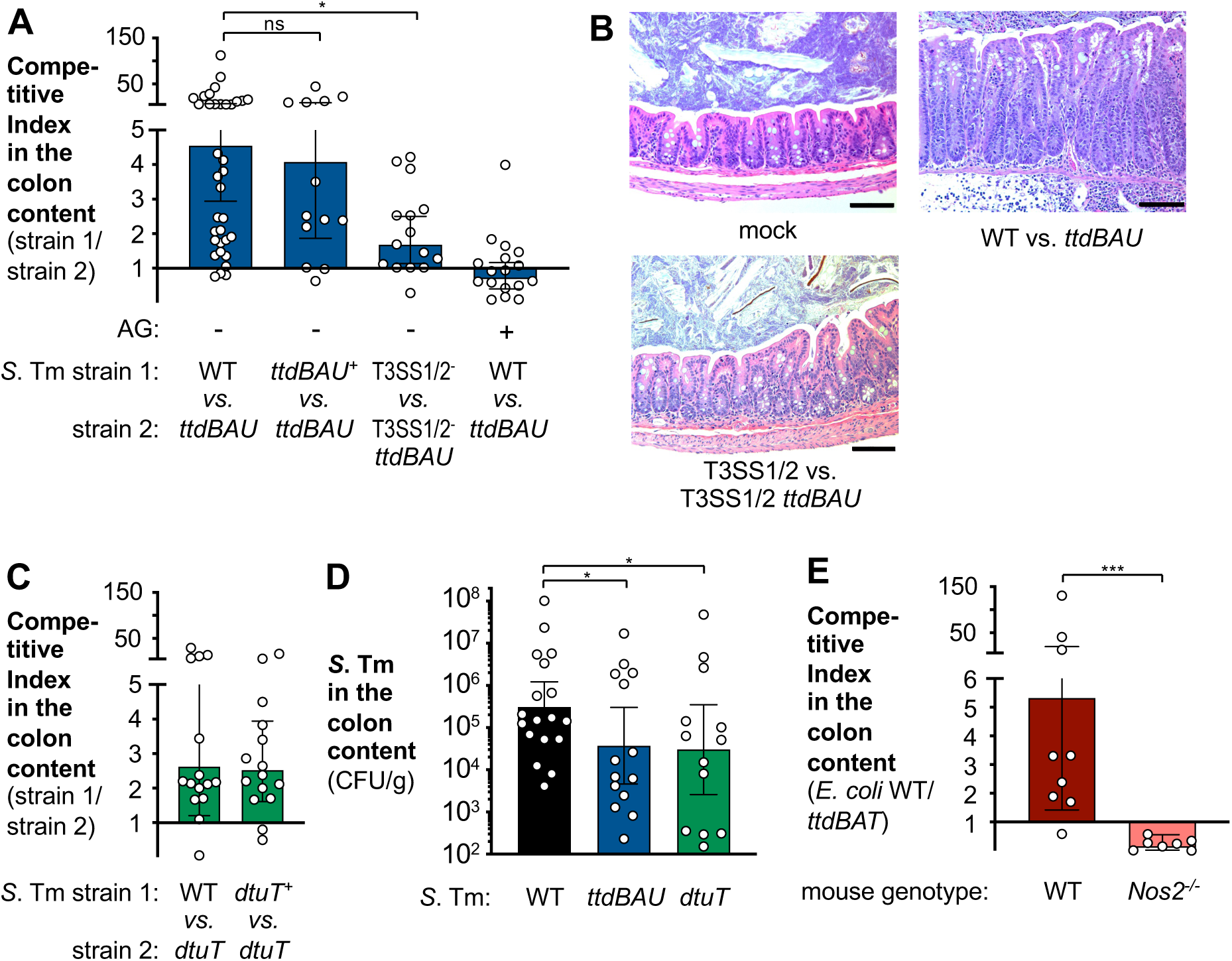
Tartrate utilization enhances fitness of Enterobacteriaceae family members during intestinal inflammation. **(A and B)** CBA mice were intragastrically inoculated with an equal mixture of the indicated *S.* Tm strains. Animals received normal water or aminoguanidine chloride (AG) in the drinking water for the duration of the experiment, as indicated. **(A)** *S.* Tm populations in the colonic content were analyzed four days after inoculation by plating on selective indicator media. The competitive index was calculated by determining the ratio of strain 1 and 2 in the colon content, normalized by the ratio of the two strains in the inoculum. **(B)** Representative images of haematoxylin and eosin-stained colonic section. **(C)** CBA mice were infected as described above with an equal mixture of the indicated *S.* Tm strains, and the competitive index in the colon content determined four days after infection. **(D)** Groups of CBA mice were inoculated with either the *S*. Tm wild-type strain, a *ttdBAU* mutant, or a *tduT* mutant. Abundance of *S*. Tm in the colon content was determined four days after infection. **(E)** C57BL/6 mice and iNOS (*Nos2*)-deficient mice on the C57BL/6 background were treated with 2 % dextran sulfate sodium (DSS) in the drinking water for four days. The animals were then intragastrically inoculated with an equal mixture of NRG857c wild-type and a *ttdBAT* mutant. Five days later, the competitive index in the colon content was determined by plating on selective indicator media. Each dot represents data from one animal. Bars represent the geometric mean +/- 95% geometric confidence interval; * *P* < 0.05; * * * *P* < 0.001; ns, not statistically significant.

### Tartrate levels increase during *S.* Tm infection

We next investigated the origin of tartrate in the murine gut. We first quantified tartrate concentrations in the gut lumen using gas chromatography-mass spectrometry GC-MS) (**Fig. 3A**). In mock-treated animals, tartrate levels were at the limit of quantification, while tartrate concentrations increased significantly during *S.* Tm infection (**Fig. 3A**). Consistent with the idea that *S*. Tm consumes tartrate, abolishing the ability to utilize tartrate (*ttdBAU* and *dtuT* mutants) further increased concentrations of free tartrate in the extracellular environment (**Fig. 3A**). Curiously, the concentration of tartrate in the diet was substantially lower than the concentration observed during *S*. Tm infection (**Fig. S3A**), which prompted us to consider non-dietary sources of tartrate.

**Figure 3:**
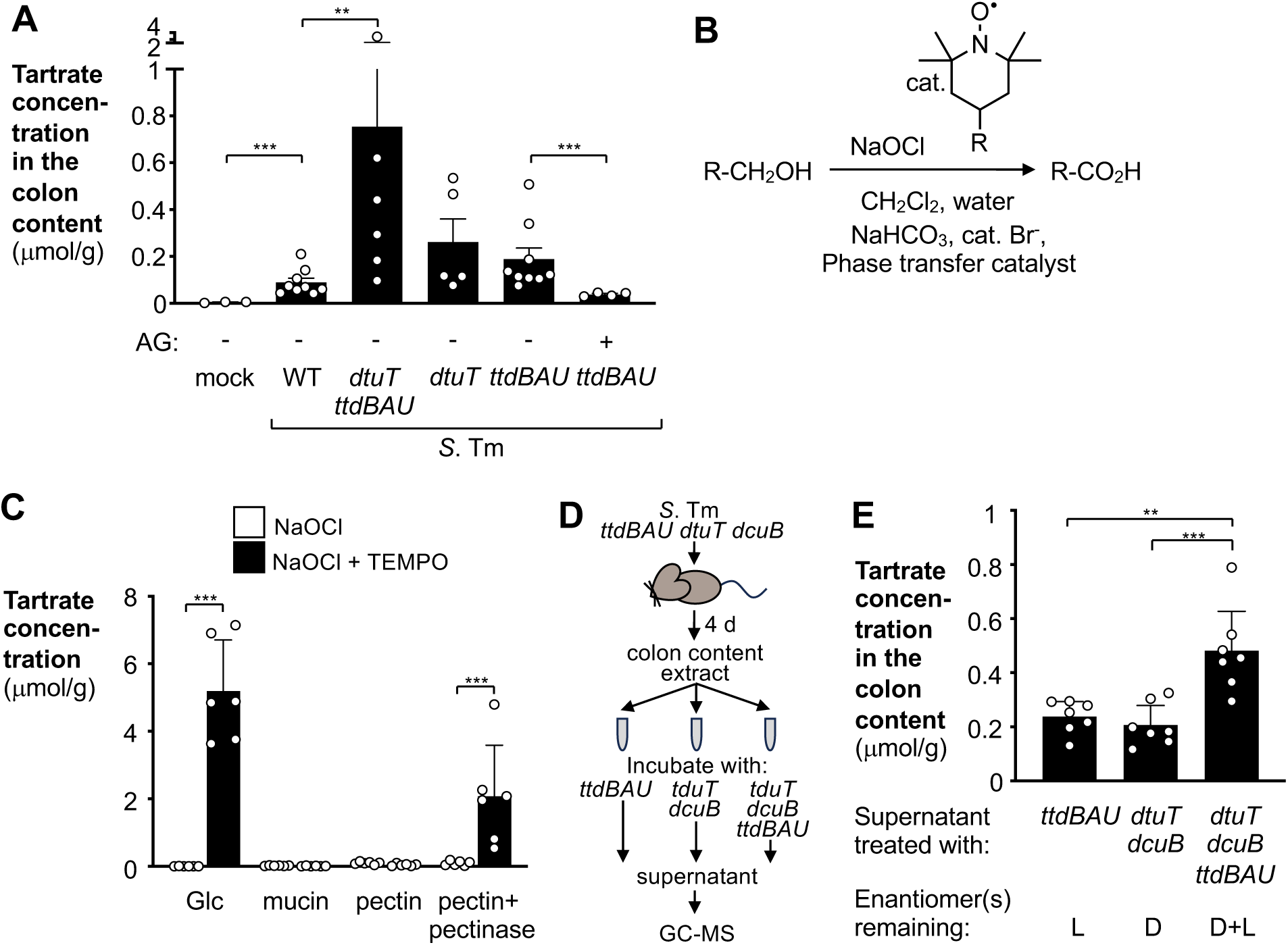
D- and L-tartrate are generated as byproducts of sugar oxidation. **A)** CBA mice were mock treated or inoculated intragastrically with the indicated *S.* Tm strains. A group of mice received aminoguanidine chloride (AG) in drinking water for four days. Tartrate concentration in the colonic content was measured by GC-MS four days after inoculation. Each dot represents data from one animal. **B)** Anelli-Montanari oxidation of primary alcohols to the corresponding carboxylic acid. **C)** Glucose, mucin, pectin, and digested pectin were oxidized with hypochlorite (NaOCl) in the absence or presence of the nitroxyl radical TEMPO. Tartrate concentration was quantified at the end of the reaction by GC-MS. Each dot represents one biological replicate. (**D and E**) Detection of D- and L-tartrate *in vivo*. CBA mice were intragastrically infected with the *S. Tm ttdBAU dtuT dcuB* mutant. After four days, the colon content was collected. Extracts of colon content were incubated in the presence of *S*. Tm mutants that are able to degrade one or both tartrate enantiomer. Tartrate in the supernatant was quantitated by GC-MS. **(D)** Schematic representation of the experimental design **(E)** Concentrations of tartrate enantiomers remaining after stereospecific degradation. Each dot represents pooled data from three animals. Bars represent the geometric mean +/- geometric standard deviation. * * *P* < 0.01, * * * *P* < 0.001.

During S. Tm infection, inflammatory reactive oxygen and nitrogen species diffuse into the gut lumen, giving rise to oxidized organic and inorganic compounds that in turn are utilized by *S*. Tm as carbon or energy sources ^24,32,33^. We therefore hypothesized that tartrate might be generated as a result of *S*. Tm-induced inflammation (**Fig. 2B**). To test this idea, we assessed whether L-tartrate utilization provides a fitness advantage in the absence of inflammation (**Fig. 2A and B**). *S*. Tm expresses two type three secretion systems to invade non-phagocytic epithelial cells (T3SS-1) and to mediate replication in the host cells and the intestinal mucosa (T3SS-2). Mutants lacking both type three secretion systems do not induce inflammation and therefore colonize the gut lumen to a lesser extent than the wild-type strain (**Fig. S3B and C**) ^34,35^. In the absence of inflammation, the T3SS-1/2 mutant and the T3SS-1/2 *ttdBAU* mutant were recovered in equal numbers (**Fig. 2A**), suggesting that tartrate availability is low in the absence of gut inflammation.

### Tartrate is a side product of Anelli’s oxidation of monosaccharides

Given that inflammation appeared to be linked to tartrate production, we examined whether inflammation-mediated, non-enzymatic oxidation of molecules in the gut plays a role in tartrate generation. Primary alcohols can be converted to the corresponding aldehyde or carboxylic acid through Anelli’s reaction, using sodium hypochlorite as a stoichiometric oxidant and stable nitroxyl radicals, such as 2,2,6,6-tetramethylpiperidine-1-oxyl (TEMPO), as catalyst (**Fig. 3B**) ^36^. As such, oxidation of glucose with hypochlorite and TEMPO is expected to yield glucaric acid ^37^. However, under suboptimal conditions, side reactions produce various breakdown products, including tartrate (**Fig. 3C**). Most carbohydrates in the large intestine are complex glycans, which are broken down by commensal microbes through diverse glycan capture and depolymerization machinery at the cell surface to yield mono- and disaccharides ^1,38^. To mimic this situation, we attempted to oxidize hog mucin and pectin with hypochlorite and TEMPO, however, little tartrate was produced in these reactions (**Fig. 3C**). We only detected tartrate when we oxidized enzymatically depolymerized pectin with hypochlorite and TEMPO (**Fig. 3C**), demonstrating that tartrate is generated through the oxidation of monosaccharides by hypochlorite and nitroxyl radicals.

### Reactive nitrogen species are required to generate tartrate during *S.* Tm infection

Reactive nitrogen (RNS) and oxygen (ROS) species, such as nitric oxide and hypochlorite, are important components of host inflammation. We hypothesized that, akin to Anelli’s oxidation *in vitro*, inflammatory RNS and ROS would oxidize sugars to form tartrate during *S*. Tm infection. The host enzyme inducible nitric oxide synthase (iNOS) generates most of the nitric oxide during inflammation ^39^. We therefore administered aminoguanidine chloride, an iNOS inhibitor, in the drinking water of *S*. Tm-infected CBA mice and quantified tartrate concentrations in the lumen (**Fig. 3A**). Consistent with our hypothesis, inhibition of iNOS significantly decreased the amount of tartrate (**Fig. 3A**). Accordingly, aminoguanidine chloride treatment abolished the fitness advantage conferred by L-tartrate utilization (*ttdBAU* mutant) (**Fig. 2A**).

Under laboratory conditions, the ratio of the different tartrate isomers generated from hexoses depends on the reaction conditions and the configuration of carbon atoms in the starting material ^40^. To determine the ratio of the two tartrate enantiomers during *Salmonella* infection, we developed an assay that relies on the stereospecific degradation of one enantiomer, and detection of the remaining enantiomer by GC-MS (**Fig. 3D and E, Fig. S4**). A group of mice was infected for four days with a *S*. Tm mutant unable to degrade D- and L-tartrate. Colon content extracts from individual mice were aliquoted, incubated with *S*. Tm mutants unable to degrade L-tartrate, D-tartrate, or both isomers (*ttdBAU dtuT dcuB* mutant), and then analyzed by GC-MS (**Fig. 3E**). The tartrate concentration in samples treated with the *ttdBAU dtuT dcuB* mutant was similar to untreated samples (**Fig. 3A**). In contrast, incubation with the *ttdBAU* and the *dtuT dcuB* mutants equally reduced tartrate levels (0.24 μmol/g and 0.20 μmol/g, respectively)(**Fig. 3E**), indicating that the concentration of both enantiomers is approximately equal (racemic mixture).

Based on our experimentation with *S.* Tm, we hypothesized that tartrate could also be generated in other settings of colitis, such as non-infectious colitis. To test this hypothesis, we investigated the tartrate metabolism of Adherent Invasive *E. coli* in a chemically-induced colitis model (dextran sulfate sodium [DSS]-induced colitis). In this mouse model, populations of *E. coli* and other commensal Enterobacteriaceae members expand as a result of the inflammatory response ^41^. The Adherent Invasive *E. coli* wild-type strain (NRG857c) outcompeted the isogenic *ttdABT* mutant (**Fig. 1H**) in the presence of L-tartrate, but not D-tartrate, confirming the stereospecific utilization of L-tartrate by this *E. coli* strain. Importantly, the Adherent Invasive *E. coli* wild-type strain displayed increased fitness in the colon lumen of DSS-treated mice (**Fig. 2E**), and this fitness advantage was negated in iNOS-deficient (*Nos2*^-/-^) mice (**Fig. 2E**). Taken together, our experiments suggest that oxidation of sugars by inflammatory reactive nitrogen and oxygen species generates tartrate isomers, including D-tartrate.

### Tartrate supports growth of *S*. Tm through fumarate respiration and the sodium pump oxaloacetate decarboxylase

We next investigated tartrate metabolism in *S*. Tm. The L-tartrate dehydratase TtdAB converts L-tartrate to oxaloacetate ^27^, which can then enter the tricarboxylic acid (TCA) cycle. Electron acceptors, released locally by professional phagocytes, allow *S*. Tm to perform an oxidative central metabolism in the inflamed gut ^31^. Curiously, supplementing culture media with tetrathionate and nitrate did not enhance fitness, and instead abolished L-tartrate utilization by *S*. Tm (**Fig. 4A**), suggesting that tartrate is not a substrate for *S*. Tm’s oxidative central metabolism.

**Figure 4:**
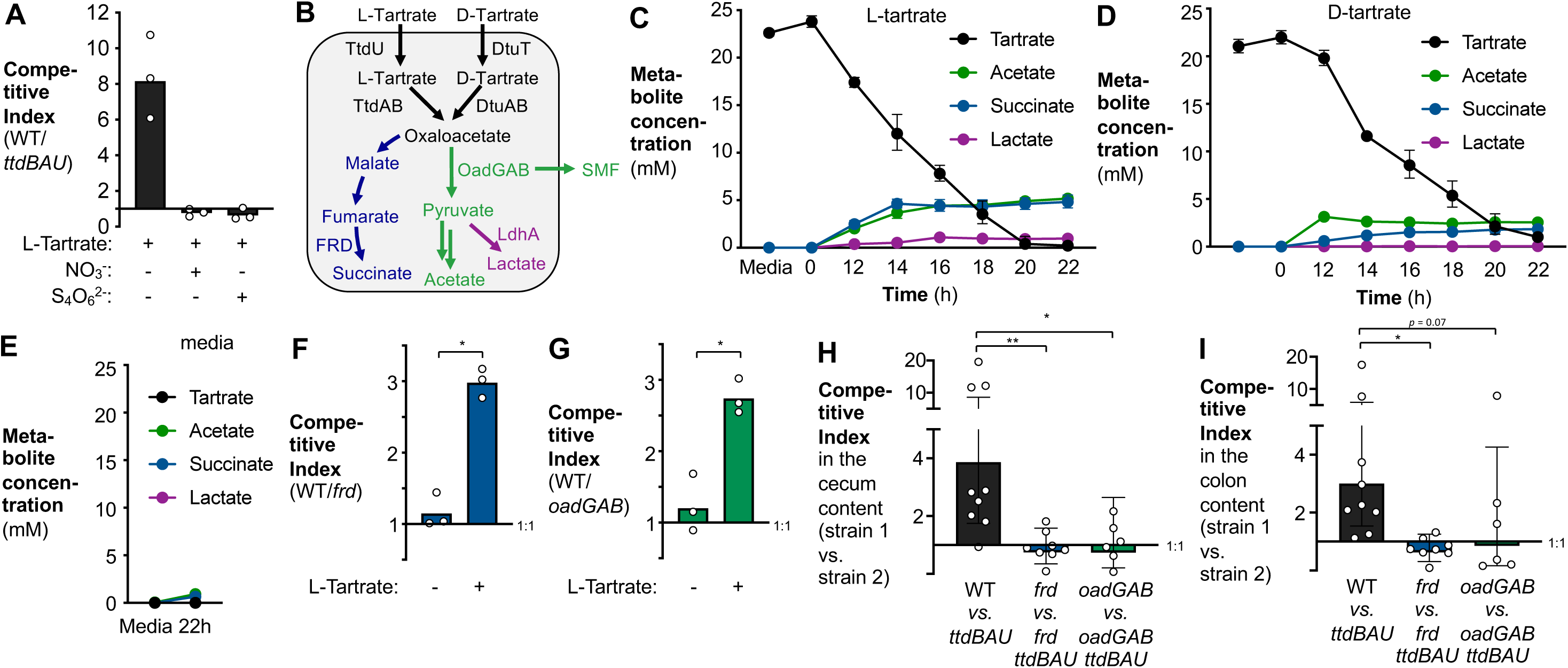
Tartrate supports fumarate respiration as well as acetate production in *S*. Tm. (**A**) Anaerobic growth of the *S.* Tm wild-type strain and a *ttdBAU* mutant in modified M9 medium supplemented with L-tartrate (20 mM), nitrate (40 mM) and tetrathionate (40 mM), as indicated. Each dot represents one biological replicate. **(B)** Schematic representation of tartrate metabolism in S. Tm, leading to succinate, acetate, and lactate production. SMF, sodium motive force. **(C-E)** The *S*. Tm wild-type strain was cultured anaerobically in the presence of L-tartrate (20 mM) (**C**), D-tartarte (20mM) (**D**), or no tartrate (**E**). Samples were collected at the indicated time points and fermentation products in the supernatant were quantitated by GC-MS. *N* = 3 biological replicates for each condition. Bars show the geometric mean +/- geometric standard error. **(F-G)** Competitive anaerobic growth in modified M9 medium in the absence or presence of L-tartrate of the *S.* Tm wild-type strain and a *frd* mutant (**F**) or a *oadGAB* mutant (**G).** Each dot represents one biological replicate. **(H-I)** CBA mice were inoculated intragastrically with an equal mixture of the indicated *S.* Tm strains. *S.* Tm populations in the cecal content (**H**) and in the colonic content (**I**) were analyzed four days after inoculation. Each dot represents one animal. Bars represent the geometric mean +/- 95% confidence interval.; * *P* < 0.05; * * *P* < 0.01.

*E. coli* converts tartrate to oxaloacetate and then to malate, which supports fumarate respiration in a bifurcated TCA cycle (**Fig. 4B**). In contrast, tartrate metabolism in *S*. Tm appears to involve oxaloacetate decarboxylase activity ^42^ (**Fig. 4B**). To investigate tartrate metabolism in *S*. Tm, we cultured the wild-type strain in the presence of L-tartrate and D-tartrate under anaerobic conditions and quantitated potential end products of fumarate respiration and anaerobic pyruvate metabolism (**Fig. 4B**). L- and D-tartrate degradation resulted in accumulation of succinate, acetate, and lactate in the supernatant (**Fig. 4C-E**), suggesting that tartrate supports both fumarate respiration and the oxaloacetate decarboxylase sodium pump ^43,44^. Mutants lacking fumarate reductase (*frdABCD*, *frd*) and oxaloacetate decarboxylase (*oadGAB*) activity displayed a fitness disadvantage during growth on L-tartrate in the laboratory (**Fig. 4F and G**). Importantly, the fitness advantage conferred by L-tartrate utilization in the murine gut was abolished in the absence of fumarate reductase and oxaloacetate decarboxylase activity (**Fig. 4H and I**), suggesting that *S*. Tm relies on both pathways for optimal tartrate degradation during infection.

### Over-oxidation of glucose by ROS and RNS produces an array of α-hydroxy carboxylic acids

Our previous results indicate that tartrate isomers are generated as a byproduct of ROS and RNS metabolism. To determine more broadly how ROS and RNS react with common biomolecules such as monosaccharides, we oxidized glucose with sodium hypochlorite in the presence and absence of TEMPO and performed GC-MS analysis to determine the end products of this oxidation reaction (**Fig. 5A and B**). Tentative assignments of features of interest suggested that glucose is degraded to various α-hydroxy acids (**Fig. 5B and C**), likely accompanied by the release of CO_2_. Quantitative, targeted GC-MS analysis revealed that glycolate, oxalate, glycerate, tartrate, and tartronate are generated by the oxidation of glucose with hypochlorite and TEMPO (**Fig. 5D – H**). Some compounds were readily generated by the oxidation with hypochlorite alone, such as glycolate and glycerate (**Fig. 5D and F**), while others, such as tartrate and tartronate (**Fig. 5 G and H**), required both hypochlorite and TEMPO. This experiment implicates inflammatory ROS and RNS in generating an array of α-hydroxycarboxylic acids through over-oxidation of monosaccharides.

**Figure 5:**
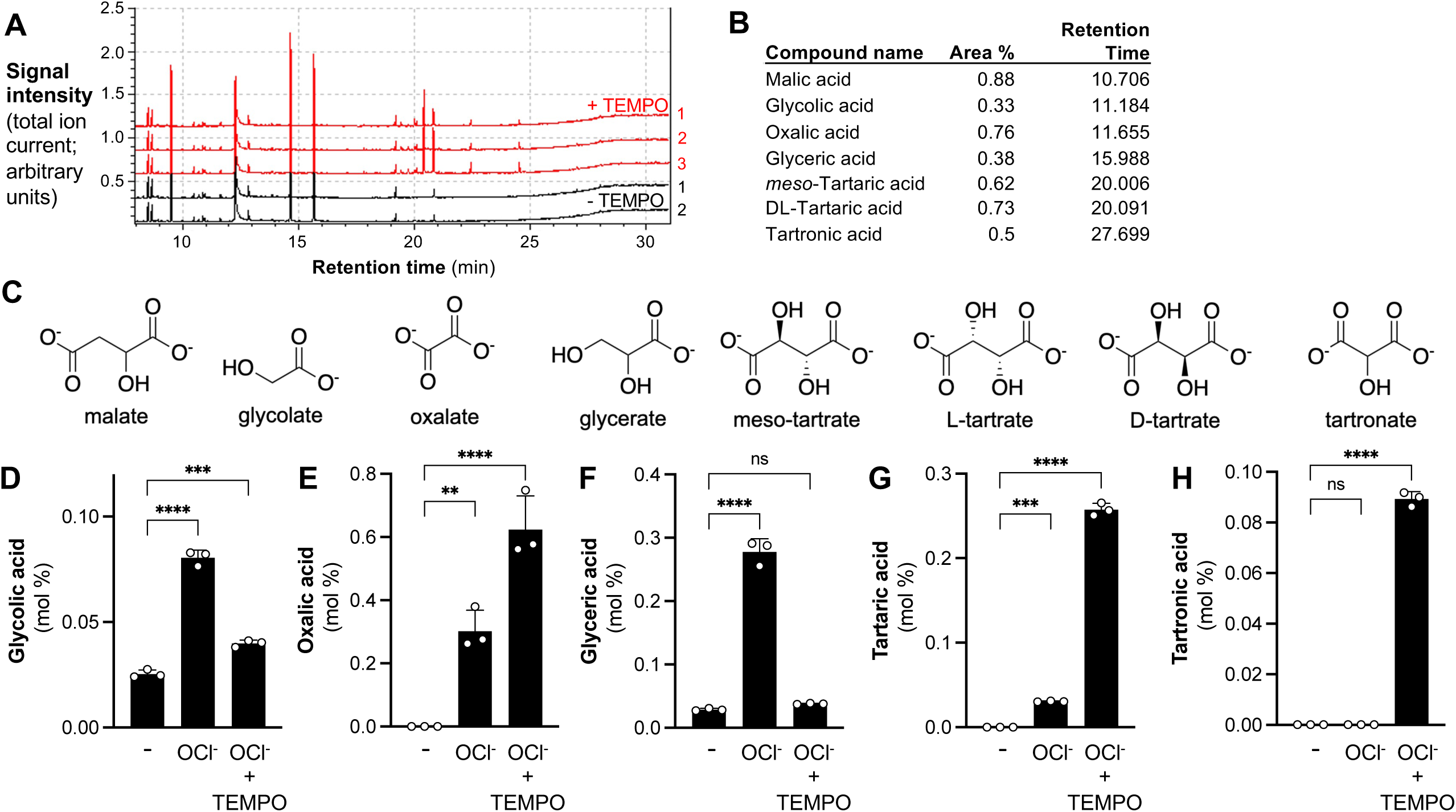
Overoxidation of glucose gives rise to various alpha-hdyroxy carboxylic acids. **(A-C)** Glucose was oxidized with hypochlorite (NaOCl) in the absence or presence of the nitroxyl radical TEMPO, and the products analyzed by GC-MS. **(A)** Chromatogram of samples treated with (red) and without (black) TEMPO. **(B)** Tentative assignments of peaks of interest. **(C)** Structural formula of compounds of interest. **(D-H)** Glucose was oxidized with hypochlorite (NaOCl) in the absence or presence of the nitroxyl radical TEMPO, and amount of glycolic acid **(D)**, oxaclic acid **(E),** glyceric acid **(F),** tartaric acid **(G),** and tartronic acid **(H)** determined using targeted GC-MS assays. Each dot represents one independent experiment. Bars show the geometric mean +/- geometric standard error.

## DISCUSSION

Tartaric acid, a C-4 dicarboxylic acid, was a key molecule in the history of molecular chirality ^45,46^. Louis Pasteur analyzed tartaric acid obtained from grape juice and compared its properties to chemically-synthesized tartaric acid, then known as paratartaric acid ^47^. Pasteur’s findings led to the discovery that paratartaric acid was a racemic mixture and composed of two tartaric acid isomers with distinct, chiral atomic structures, now denoted as L-(+)-tartrate [(2R,3R)-tartrate; *d*-tartrate in the older literature] and D-(-)-tartrate [(2S,3S)-tartrate]. To further support his conclusions, Pasteur developed a method to produce D- and L-tartrate from the racemic mixture through the stereoselective degradation by microbes ^48,49^. He recognized that biological samples would solely contain L-tartrate, however, he was unable to deduce a natural source of D-tartrate and any biological origin remained enigmatic. To date, the only way to produce D-tartrate was through stereoselective organic synthesis. As such, D-tartrate could be classified as a xenobiotic ^50^. Here, we report that both L- and D-tartrate isomers are produced during gut inflammation through RNS-catalyzed oxidation of sugars by ROS. In contrast to the production of L-tartrate by the Smirnoff-Wheeler pathway, our experiments imply a natural, albeit non-enzymatic source of D-tartrate.

The production of tartrate during colitis generates a transient nutrient niche that is exploited by Enterobacteriaceae family members. The inflammatory host response elicited by non-typhoidal *Salmonella* serovars, such as *S*. Tm, alters the gut environment, enabling the pathogen to outgrow the microbiota and to enhance transmission by the fecal-oral route ^51,52^. Here, we found an example of how host-generated metabolites support *S*. Tm gut colonization early during the infection process. In the anaerobic gut lumen, *S*. Tm relies on a bifurcated TCA cycle and fumarate reduction for ecosystem invasion ^53,54^. We found that during *S*. Tm infection, D- and L-tartrate are converted to oxaloacetate by the respective dehydratases (hydro-lyases, DtuAB and TtdAB) (**Fig. 4**). This oxaloacetate is then converted to malate and fumarate, the substrate for fumarate reduction. A second fate of oxaloacetate is its conversion to pyruvate, which is further broken down to lactate and acetate. Conversion of oxaloacetate to pyruvate is catalyzed by the oxaloacetate decarboxylase (OadGAB) sodium pump, implicating sodium motive force generation in early gut colonization. At later stages of infection, S. Tm rewires its energy metabolism in response to changes in host metabolism and inflammatory cell infiltration. A change in colonocytes energy metabolism allows oxygen influx into the gut lumen, while neutrophils transmigrating into the gut lumen generate tetrathionate ^24,32,55^. The increased availability of oxygen, tetrathionate, and nitrate allow *S*. Tm to perform anaerobic respiration and support an oxidative central metabolism at later stages of gut infection ^31^. Tartrate utilization is abrogated in the presence of oxygen, tetrathionate, and nitrate (**Fig. 4A**) ^56^, suggesting that the nutrient niche provided by tartrate is transient, and utilization subsides when better electron acceptors become available.

The ability of *Salmonella* to degrade L-tartrate (Jordan’s tartrate agar) is used for the diagnostic differentiation of some pathogenic Enterobacteriaceae family members ^12,56^, and our findings serve as an explanation for the physiological basis of this clinical test. Clinical laboratories differentiate between typhoidal serovar Paratyphi B and the closely related serovar Java strains based on L-tartrate utilization, with only gastroenteritis-associated *S.* Java able to ferment L-tartrate (**Fig. S1**) ^12,13,57^. While non-typhoidal, gastrointestinal *Salmonella* enterica serovars are typically associated with acute gastroenteritis, typhoidal serovars, such as serovars Typhi and Paratyphi A, B, and C cause a febrile illness termed typhoid fever. Typhoidal *Salmonella* strains disseminate to systemic body sites and colonize the gall bladder. Bacteria are excreted back into the intestinal tract for fecal-oral transmission ^58^. The genome of typhoidal *Salmonella* and other extraintestinal serovars contains a large number of pseudogenes, suggesting that typhoidal *Salmonella* arose from a non-typhoidal, gastrointestinal ancestor and lost the ability to benefit from inflammation-derived metabolites ^59^. Accordingly, the *dtu* gene cluster is absent from *S*. Typhi and *S*. Paratyphi A genomes; in Paratyphi C and in the extraintestinal serovar Choleraesuis, the STM0764 orthologue in the *dtu* gene cluster exhibits a frame shift ^59^. This suggests that tartrate utilization might be an adaption of the metabolism of gastroenteritis-associated bacteria to the inflamed gut.

Tartrate utilization also occurs in other microbes, such as commensal Enterobacteriaceae family members ^14^. In our study, we found that a Crohn’s Disease-associated Adherent Invasive *E. coli* strain utilizes L-tartrate, and utilization of tartrate enhances fitness in a mouse model of intestinal inflammation (**Fig. 1H and 2E**). As such, the ability to degrade tartrate might contribute to the gut microbiota changes in the context of non-infectious colitis ^60^.

## ACKNOWLEDGMENTS

We thank Dr. Julie Pfeiffer (UT Southwestern Medical Center) for helpful discussion.

## Author contributions

L.S., M.G.W., M.K.M, V.K.R., R.B.C., W.Z., E.R.H., S.T., F.F. performed and analyzed the *in vivo* and *in vitro* experiments. R.L.S. and T.F.C. performed the histopathology analysis. S.P., H.A.P. and M.M.C, contributed critical tools. T.Q., contributed to GC-MS measurements. L.S., M.G.W., M.K.M, and S.E.W. designed the experiments, interpreted the data, and wrote the manuscript with contributions from all authors.

## Data and materials availability

Materials transfer agreements and regulatory permits may be required to transfer materials described in this study. All data is available in the main text or the supplementary materials.

## METHODS

### Bacterial culture

Bacterial strains were routinely grown aerobically at 37°C in Luria-Bertani (LB) broth (10 g/L tryptone, 5 g/L yeast extract, 10 g/L sodium chloride) or on LB agar plates (10 g/L tryptone, 5 g/L yeast extract, 10 g/L sodium chloride, 15 g/L agar). When necessary, antibiotic was added to the medium at the following concentration: nalidixic acid (Nal) 50 mg/L, carbenicillin (Carb) 100 mg/L, kanamycin (Kan) 50 mg/L, and chloramphenicol 15 mg/L. For competitive growth assays involving *S.* Tm and *E. coli* strains, the chromogenic substrates 5-Bromo-4-chloro-3-indolyl phosphate (X-phos) or 5-bromo-4-chloro-3-indolyl- β-D-galactopyranoside (X-gal) was used to detect the acid phosphatase (PhoN) or β galactosidase activity, respectively. In these experiments, the wild-type strain is represented by a *S*. Tm *phoN* or *E. coli lacZ* mutant; inactivation of these genes has no fitness effect in the models tested. The competitive index was calculated by dividing the number of wild-type bacteria by the number of mutant bacteria in the output, and then normalized by the same ratio obtained from the inoculum.

### Jordan’s tartrate agar

Jordan’s tartrate agar (10 g/L pancreatic digest of casein, 10 g/L tartrate, 5 g/L sodium chloride, 15 g/L agar, 0.024 g/L phenol red) was used for the detection of tartrate fermentation. Phenol red is used as indicator of acid production. Tartrate fermentation acidifies the medium causes the lower portion of the tube to turn yellow.

### Construction of plasmids

To generate pLS9, pLS10, pLS11, pLS12, pMW337, and pMW427 (**Table S1**), the upstream and downstream regions of *ttdBA*, *ttdU*, *ttdBAU*, *frdBACD*, *oadGAB*, and *ttdBAT* were amplified by PCR from the IR715 and NRG857c chromosomes, respectively (**Table S2**). PCR products were purified and inserted into SphI-digested pGP706 using Gibson Assembly Master Mix, as per the manufacturer’s recommendations (NEB, MA). Successful insertions of the flanking regions were verified by Sanger sequencing. All plasmids were propagated in DH5α λ*pir*. To generate pRC14 and pRC17, the predicted promoter and coding sequence of the IR715 *ttdBAU* operon and the STM0765 (*dtuT*) gene were amplified and cloned into the SphI restriction enzyme site of pSW327.

### Generation of bacterial mutants

To generate LS24, LS25, LS26, LS45, LS74, MW399, and MM54 (**Table S1**), suicide plasmids were introduced into S17-1 λ*pir* and conjugated into *S*. Tm strain IR715 or NRG857c, respectively, by bi-parental mating. The ex-conjugants (first crossover event) were selected on LB supplemented with nalidixic acid and kanamycin, or nalidixic acid and chloramphenicol, as required. Sucrose selection was utilized to select for bacteria in which a second crossover event had occured ^61^. The resulting strains were verified by PCR. The ΔS*TM0765*::Kan^R^ mutation and the ΔS*TM0762*::Kan^R^ mutation ^62^ was transduced by phage P22 HT *int-*105 ^63^ from *S.* Tm 14028 to IR715 to generate LS118 and LS115, respectively. To allow for transduction of clean, unmarked mutations, the suicide plasmid used to generate this mutation was re-introduced into the mutant background, transduced into the background of interest, and then removed using sucrose selection as described above. Using this approach, the *ttdBAU* mutation was transduced from LS25 into SPN487, LS118, LS74, MW399 and LS170 to generate LS31, LS123, LS49, LS80 and LS171 respectively. Moreover, the *dcuB* mutation was transduced from LS45 into LS118 to generate LS170. pRC14 and pRC17 were integrated into LS25 and LS118 to give rise to RC103 and RC128, respectively. Integration of pRC14 and pRC17 into the *phoN* gene was confirmed by PCR and lack of acid phosphatase activity in the resulting strain.

### Growth assays

Sterile M9 minimal medium (M9 minimal salts, 1 mM MgSO_4_, 0.1 mM CaCl_2_) ^64^, was supplemented with 0.2% casamino acids and 20 mM glycerol. Carbon sources were added to designated samples at a final concentration of 20 mM. Sodium nitrate and potassium tetrathionate were added to a final concentration of 40 mM where needed. Modified M9 media was inoculated with a 1:1 ratio of the indicated strains and incubated under anaerobic condition for 16 hours. The CFU/mL of each strain was determined by plating serial dilutions on indicator agar. Samples for GC-MS analysis were collected to quantify tartrate concentration.

### Animal models of colitis

Male and female CBA, C57BL/6 and *Nos2*-deficient (on the C57BL/6 background) mice were originally obtained from the Jackson Laboratory and bred at UT Southwestern and UC Davis. Mice were randomly assigned into cages 3 days prior to experimentation. Animals were housed in individually ventilated cages with *ad libitum* access to water and feed. At the end of the experiments, mice were humanely euthanized using carbon dioxide inhalation. All mouse experiments were performed in accordance with the Institutional Animal Care and Use Committee at UT Southwestern Medical Center and UC Davis.

#### CBA colitis model

Groups of 8-10 weeks old CBA mice were intragastrically inoculated with 1 x 10^9^ CFU for single strain infection experiments, 5 x 10^8^ CFU of each strain for competitive infection experiments, or mock treated (LB broth). After 4 days, mice were euthanized and colonic content, cecal content, and tissue collected. To evaluate the contribution of iNOS activity, groups of mice were treated with 1 mg/mL aminoguanidine chloride in their drinking water for all 4 days of the experiment, as indicated. Colonic and cecal contents were serially diluted on selective agar plates to determine CFU/g of each *S.* Tm strain.

#### Chemically-induced colitis model

Groups of C57BL/6 wild-type and *Nos2*-deficient mice received 2 % dextran sulfate sodium (DSS; FisherScientific) in the drinking water. At the onset of inflammation ^25^, four days after begin of the DSS treatment, an equal mixture of the NRG857c *lacZ* mutant and the isogenic *ttdABT* mutant (5 x 10^8^ CFU each strain) were administered by gavage and DSS treatment continued for another 4 days. Animals were switched to regular water one day before euthanasia. Samples of the cecal and colon content were collected in PBS, diluted in PBS, and spread on selective chromogenic agar plates to determine CFU/g of each *E. coli* strain.

### Histopathology analysis

Cecal and colonic tissues were fixed in 10% phosphate-buffered formalin for 48 h and stored in 70% ethanol. Samples were embedded in paraffin, and 5 μm sections were stained with hematoxylin and eosin. Stained sections were blinded and scored ^31^.

### *In vitro* sugar oxidation

Sugar oxidations were performed at room temperature and pH was kept constant at 8 ^33^. Either 0.4 g of glucose or mucin was dissolved in 50 mL of water. Pectin was depolymerized with 100 U of pectinase in 0.2 M acetate buffer (pH 4.5) for 3 hours at 50 °C. TEMPO (2,2,6,6-tertamethyl-1-piperidyloxy) was added to the solution, as indicated. Sodium hypochlorite, 2.2 ml, was added to the reaction and the pH was kept constant at 8 for the entire reaction. The reaction was then precipitate with 95% ethanol and incubated for 48 hours 4°C. After a centrifugation at 2,000 g for 5 min at 4°C, the pellet was washed with 20 mL of acetone. Samples were then centrifuged as described above. The resulting powder was dried overnight and analyzed by GC-MS.

### Gas chromatography-mass spectrometry

For mouse experiments, colon and cecal content from mice was collected in sterile PBS and samples were vortexed for 2 min and centrifuged at 6,000 g for 15 min at 4 °C. For *in vitro* bacterial cultures, 2 mL of culture were centrifuged at 20,000 g at 4 °C. For samples derived from TEMPO reactions, 20 mg of the resulted powder was resuspended in 1 mL PBS. All samples were aliquoted in 0.1 mL portions and succinic-2,2,3,3,-d_4_ acid was added as an internal standard. Samples were vacuum centrifuged for 2 h and stored at -80 °C. Samples were resuspended in 50 μL 20 mg/mL methoxyamine-hydrochloride in dry pyridine, individually sonicated for 1 min and incubated for 20 min at 80°C. Then, 0.1 mL of *N*-*tert*- Butyldimethylsilyl-*N*-methyltrifluoroacetamide with 1 % *tert*-Butyldimethylchlorosilane was added, and the samples incubated for 1 h at 80°C. Samples were centrifuged for 1 minute at 16,000 x g and the supernatant was transferred to autosampler vials for gas chromatography-mass spectrometry (GC-MS) analysis (Shimadzu, TQ8040). The injection temperature was 250°C and the injection split ratio was set to 1:100 with an injection volume of 1 μL. The oven temperature started at 50 °C for 2 min, increasing to 100 °C at a rate of 20 °C per min with a holding temperature of 3 min, and to 330 °C at a rate of 40 °C per min with a final hold at this temperature for 3 min. Flow rate of the helium carrier gas was kept constant at a linear velocity of 50 cm/s. The column used was a 30 m × 0.25 mm × 0.25 μm Rxi-5Sil MS (RESTEK). The interface temperature was 300°C. The electron impact ion source temperature was 200 °C, with 70 V ionization voltage and 150 μA current. For qualitative experiments, Q3 scans (range of 50-550 m/z, 1000 m/z per second) were performed. Subsequent quantitative measurements were performed using pure, commercially available compounds as external standards. The mass spectrometry method, monitored m/z, and quantifier (in italics) are as follows: Acetate (SIM; *117*, 75, 159), lactate (MRM; *261*>*233,;* 261>189), succinate (MRM; *289>147*, 331>189), deuterated succinate (MRM; *293>147*, 335>189), tartrate (MRM; *389>147*, 549>417), glycolic acid (SIM; *247*, 219, 163), glyceric acid (SIM; *391*, 231, 115), oxalic acid (SIM; *261*, 189, 147), tartronic acid (SIM; *405*, 377, 263). Recovery of target metabolites was determined using deuterated compounds as internal standards. Quantification was based on external standards comprised of a series of dilutions of pure compounds, derivatized as described above at the same time as the samples.

To differentiate between D- and L-tartrate, samples were pre-digested using bacterial strains with defined lesions in D- and L-tartrate utilization. For in vitro measurements, Jordan’s Tartrate Agar base was supplemented with either 1 mM D-tartrate, 1 mM L-tartrate, or 0.5 mM of each isomer. Samples with a volume of 0.3 mL were inoculated with 0.015 mL of various *S*. Tm overnight cultures, as indicated, and incubated at 37 °C overnight. Bacterial cells were removed by centrifugation at 20,000 g at 4°C. The supernatant was collected, internal standard added, and 0.1 mL aliquots processed for GC-MS as described above. For *in vivo* measurements, fifteen CBA mice were infected with a *S*. Tm *dtuT ttdBAU dcuB* mutant for 4 days as described above. The colon content was suspended in 2 ml of PBS by vortexing. The suspension was centrifuged at 20,000 g for 15 min at 4°C to remove particles and bacterial cells. The supernatants from three animals were combined and filter sterilized. Aliquots of 0.3 mL were inoculated with 0.015 ml of various *S*. Tm overnight cultures, as indicated, and incubated at 37 °C overnight. Bacterial cells were removed by centrifugation at 20,000 g at 4°C. The supernatant was collected, internal standard added, and 0.1 mL aliquots processed for GC-MS as described above.

### Statistical analysis

Data analysis was performed in GraphPad Prism v8.4.1. Values of bacterial population sizes, competitive indices, were normally distributed after transformation by the natural logarithm. A two-tailed Student’s *t*-test was used for ln-transformed data. Unless otherwise stated, *, *P* < 0.05; **, *P* < 0.01; ***, *P* < 0.001; ns, not statistically significant. In all mouse experiments, *n* refers to the number of animals from which samples were taken. Sample sizes (i.e. the number of animals per group) were not estimated a priori since effect sizes in our system cannot be predicted. No predicted statistical outliers were removed since the presence or absence of these potential statistical outliers did not affect the overall interpretation. Mice that were euthanized early due to health concerns were excluded from analysis.

