## Supplementary Figures for "Byproducts of inflammatory radical metabolism provide transient nutrient niches for microbes in the inflamed gut"

**A**

| Organism: | D-tartrate | L-tartrate |
|---|---|---|
| *E. coli* (K-12) | + | + |
| *E. coli* EHEC (86-24) | + | + |
| *E. coli* EPEC (2348/69) | + | + |
| *C. rodentium* (DBS100) | + | - |
| *K. oxytoca* (M5al) | - | + |
| *Salmonella* Typhimurium (14028) | + | + |
| *Salmonella* Typhimurium (SR11) | + | + |
| *Salmonella* Typhimurium (SL1344) | + | + |
| *Salmonella* Typhimurium (14028) | + | + |
| *Salmonella* Enteritidis (MZ0630) | - | + |
| *Salmonella* Newport (MZ1838) | + | + |
| *Salmonella* Typhi (Ty2) | - | + |
| *Salmonella* Paratyphi A (ATCC9150) | - | - |
| *Salmonella* Paratyphi B (DMS2471) | - | - |
| *Salmonella* Paratyphi B (DMS155) | - | - |
| *Salmonella* Paratyphi B *var.* Java (DMS106) | - | + |
| *Salmonella* Paratyphi B *var.* Java (IP83) | - | - |
| *Salmonella* Paratyphi C (IP33K) | - | + |
| *Y. enterocolitica* (WA314) | + | + |

**B**

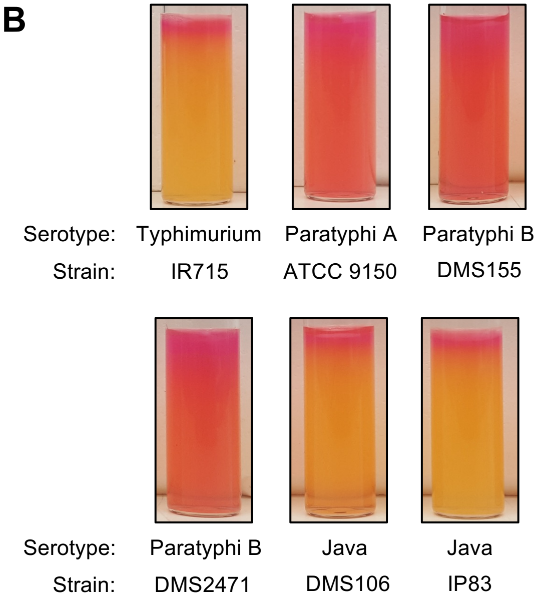


**Figure S1: Utilization of D- and L-tartrate by gut bacteria.**

**(A)** Jordan's tartrate agar base, supplemented with either D- or L-tartrate, was stab-inoculated with the indicated bacterial strains. Tubes were incubated at 37ºC overnight. A bright yellow color is indicative of tartrate fermentation (shown as +; absence of a color change is denoted as -).

**(B)** Representative images of L-tartrate utilization by typhoidal and non-typhoidal *Salmonella* serovars.

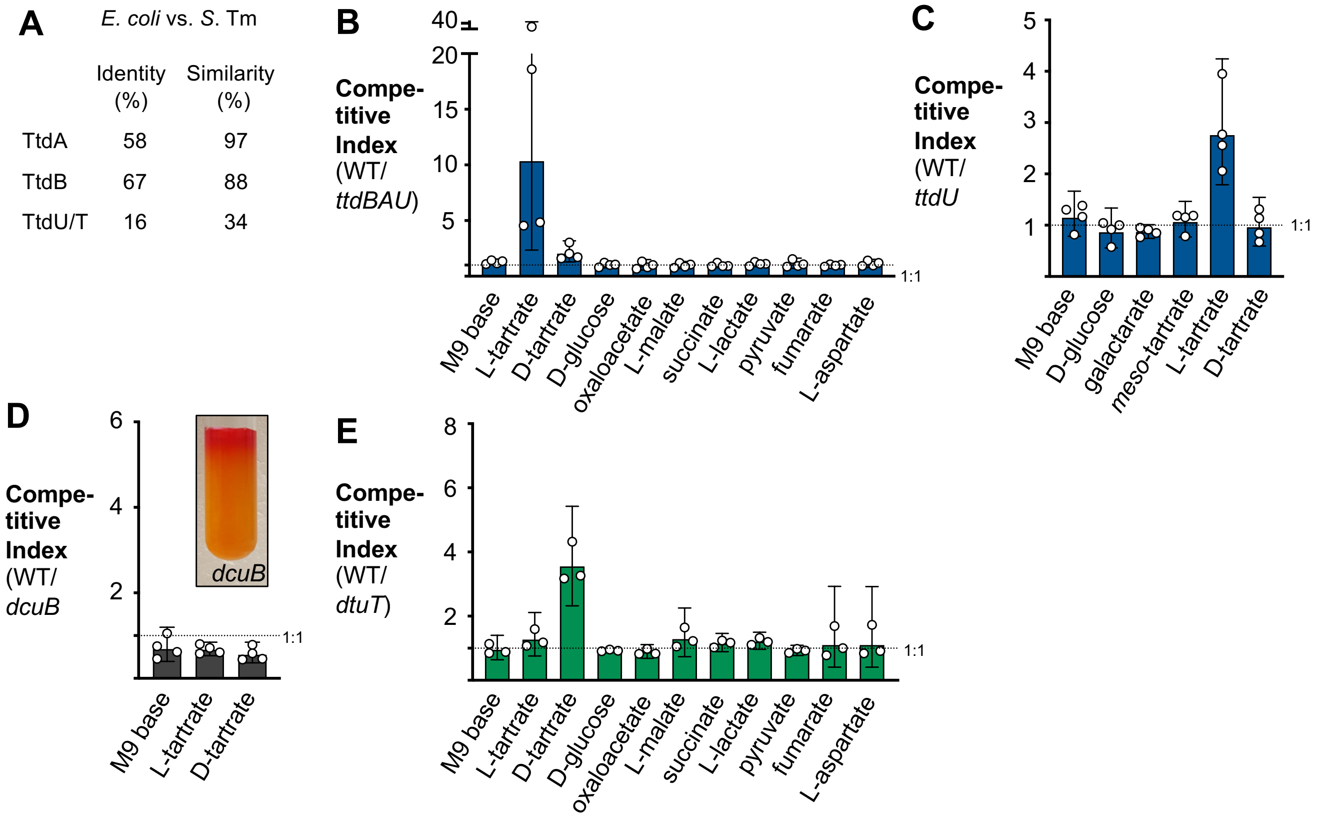


| | Identity (%) | Similarity (%) |
|---|---|---|
| TtdA | 58 | 97 |
| TtdB | 67 | 88 |
| TtdU/T | 16 | 34 |

**Figure S2: Genetic requirements for the utilization of tartaric acid by *Salmonella* Typhimurium.**
**(A)** Comparison of the primary amino acid sequence of proteins involved in L-tartrate utilization.
**(B-C)** Competitive anaerobic growth in modified M9 media of the *S.* Tm wild-type strain and a Δ*ttdBAU* mutant (**B**) or a a Δ*ttdU* mutant (**C**). The competitive index was determined 16 hr after growth under anaerobic conditions in the absence or presence of different carbon sources (20 mM).
**(D)** Competitive anaerobic growth in modified M9 media in the absence or presence of L-tartrate (20 mM) and D-tartrate (20 mM) of the *S.* Tm wild-type and a *dcuB* mutant. The competitive index was determined after 16 hr of anaerobic growth. The insert shows a *dcuB* mutant grown in Jordan's tartrate agar base containing D-tartrate.
**(E)** Competitive anaerobic growth in modified M9 media of the *S.* Tm wild-type strain and a Δ*dtuT* mutant. The competitive index was determined 16 hr after growth under anaerobic conditions in the absence or presence of different carbon sources (20 mM).
Each dot represents one biological replicate. Data are shown as geometric mean +/- geometric standard deviation.

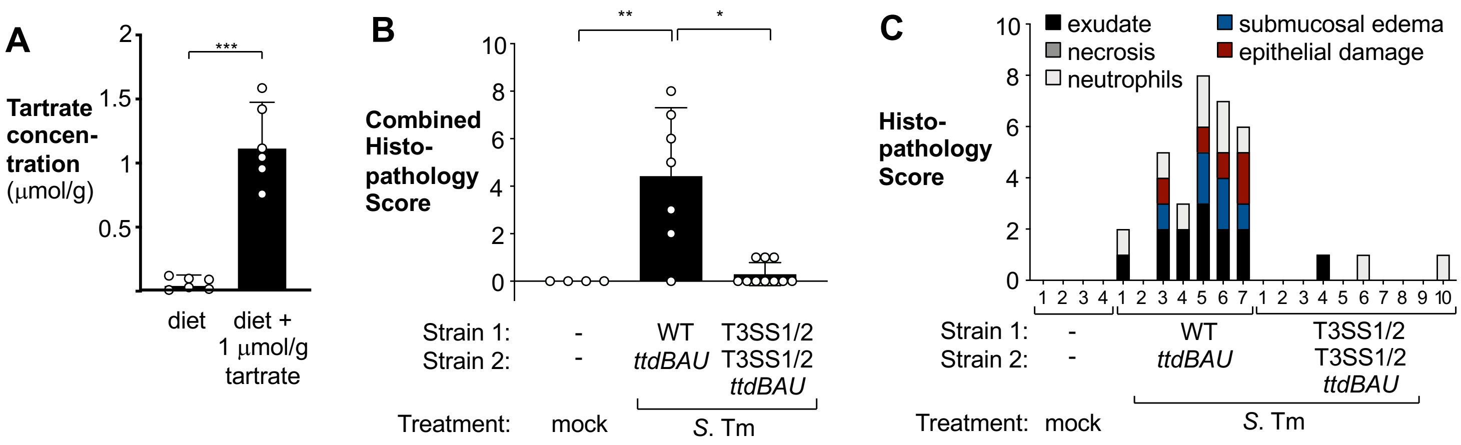


**Figure S3: Tartrate availability and gut inflammation.**
**(A)** Tartrate concentration in the mouse diet was measured by GC-MS with or without the experimental addition of L-tartrate. Each dot represents one independent measurement.
**(B-C)** CBA mice were mock treated or inoculated intragastrically with an equal mixture of the indicated *S*. Tm strains (see **Fig. 2A and B**). Colonic sections were analyzed four days after infection. Combined histopathology score of pathological lesions **(B)** and semi-quantitative histopathology score for each animal.
* $P < 0.05$; * * $P < 0.01$.

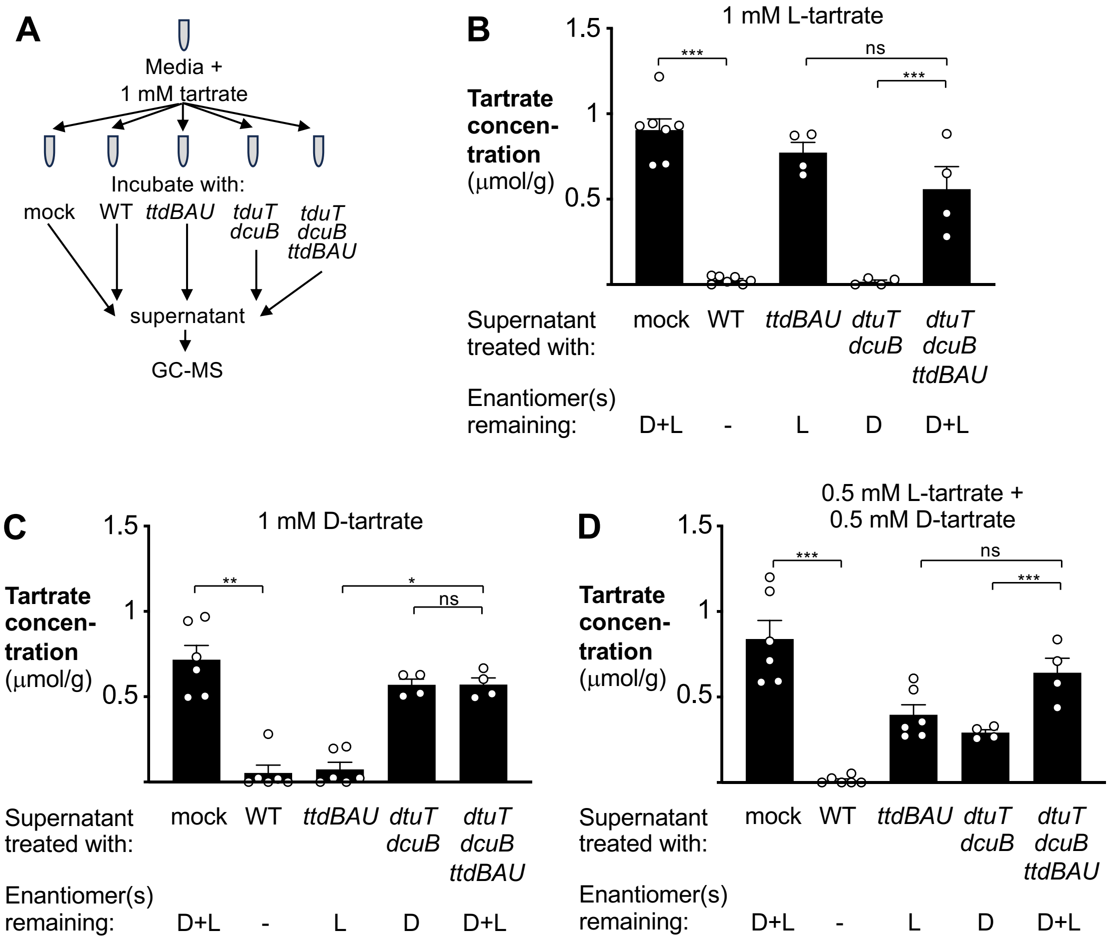


**Figure S4: Quantitation of D- and L-tartrate under defined laboratory conditions.**

**(A-D)** Jordan's tartrate agar base was supplemented with tartrate enantiomers. Samples were inoculated with *S*. Tm strains that differed in their ability to degrade tartrate enantiomers. The remaining tartrate in the supernatant was quantitated by GC-MS.

**(A)** Schematic representation of the experimental design.

**(B)** Media was supplemented with 1 mM L-tartrate.

**(C)** Media was supplemented with 1 mM D-tartrate.

**(D)** Media was supplemented with 0.5 mM of each L- and D-tartrate

Each dot represents one biological replicate. Bars show the geometric mean +/- geometric standard deviation.
