## Supplemental Tables for "Byproducts of inflammatory radical metabolism provide transient nutrient niches for microbes in the inflamed gut"

**Table S1:** Recombinant DNA and Bacterial strains used in this study

| **Plasmid** | **Description** | **Reference** |
|---|---|---|
| pGP704 | Plasmid: *ori*(R6K) *mobRP4* Amp/Carb$^R$ | [1] |
| pSW327 | Plasmid: Internal fragment of the *phoN* coding sequence cloned into pGP704 | [2] |
| pGP706 | Plasmid: *ori*(R6K) *mobRP4 sacRB* Kan$^R$ | [3] |
| pLS9 | Plasmid: upstream and downstream region of *S*. Tm *ttdBA* cloned into pGP706 | This study |
| pLS10 | Plasmid: upstream and downstream region of *S*. Tm *ttdU* cloned into pGP706 | This study |
| pLS11 | Plasmid: upstream and downstream region of *S*. Tm *ttdBAU* cloned into pGP706 | This study |
| pLS12 | Plasmid: upstream and downstream region of *S*. Tm *frdABCD* cloned into pGP706 | This study |
| pMW337 | Plasmid: upstream and downstream region of *S*. Tm *oadGAB* cloned into pGP706 | This study |
| pSW301 | Plasmid: upstream and downstream region of *S*. Tm *dcuB* cloned into pRDH10 | [2] |
| pRC14 | Plasmid: *S*. Tm *ttdBAU* promoter and coding sequence cloned into pSW327 | This study |
| pRC17 | Plasmid: *S*. Tm *dtuT* promoter and coding sequence cloned into pSW327 | This study |
| pMW427 | Plasmid: upstream and downstream region of *E. coli ttdABT* cloned into pGP706 | This study |
| **Strain** | **Description** | **Reference** |
| DH5*α* λ*pir* | *E. coli,* F$^-$ endA1 hsdR17 (r$^-$ m$^+$) supE44 thi-1 *recA1 gyrA relA1* Δ(*lacZYA-argF)U189* *Φ*80*lacZ*ΔM15 λ*pir* | [4] |
| S17-1 λ*pir* | *E. coli,* zxx*::RP4 2-(*Tet$^R$::Mu*) (*Kan$^R$::Tn7*)* λ*pir recA1 thi pro hsdR* (r$^-$ m$^+$) | [5] |
| IR715 | *S*. Typhimurium ATCC14028 Nal$^R$ | [6] |
| AJB715 | IR715 *phoN*::Kan$^R$ | [7] |

| | | |
|---|---|---|
| SPN487 | IR715 Δ*invA*(-9 to +2057) Δ*spiB*(+25 to +1209) | [8] |
| SGD947 | 14028 ΔSTM0762::Kan$^R$ (=Δ*dtuA*::Kan$^R$) | [9] |
| SGD950 | 14028 ΔSTM0765::Kan$^R$ (=Δ*dtuT*::Kan$^R$) | [9] |
| SW1401 | IR715 Δ*invA* Δ*spiB* *phoN*::Kan$^R$ | [2] |
| LS24 | IR715 Δ*ttdBA* | This study |
| LS25 | IR715 Δ*ttdBAU* | This study |
| LS26 | IR715 Δ*ttdU* | This study |
| LS31 | IR715 Δ*invA* Δ*spiB* Δ*ttdBAU* | This study |
| LS45 | IR715 Δ*dcuB* | This study |
| LS49 | IR715 Δ*frdABCD* Δ*ttdBAU* | This study |
| LS74 | IR715 Δ*frdABCD* | This study |
| LS80 | IR715 Δ*oadGAB* Δ*ttdBAU* | This study |
| LS115 | IR715 Δ*dtuA*::Kan$^R$ | This study |
| LS118 | IR715 Δ*dtuT*::Kan$^R$ | This study |
| LS123 | IR715 Δ*dtuT*::Kan$^R$ Δ*ttdBAU* | This study |
| LS170 | IR715 Δ*dtuT*::Kan$^R$ Δ*dcuB* | This study |
| LS171 | IR715 Δ*dtuT*::Kan$^R$ Δ*dcuB* Δ*ttdBAU* | This study |
| MW399 | IR715 Δ*oadGAB* | This study |
| RC103 | IR715 Δ*ttdBAU* *phoN*::*ttdBAU* (=*ttdBAU*$^+$) | This study |
| RC128 | IR715 Δ*dtuT*::Kan$^R$ *phoN*::*dtuT* (*dtuT*$^+$) | This study |
| NRG857c | AIEC wild-type strain | [10] |
| LB33 | NRG857c Δ*lacZ* | [11] |
| MM54 | NRG857c Δ*ttdABT* | This study |

**Table S2:** Primers used in this study.

| **Targeted mutagenesis and complementation constructs** | | |
|---|---|---|
| Purpose | Sequence (restriction enzyme sites are shown in italics, sequences homologous to target genes are underlined) | Reference |
| Deletion of *ttdBA* in *S*. Tm | 5'-CTAGAGGTACCGCATGCCGGTTGTATTGGCGCTC-3'<br>5'-AGCCTTAATAAGGATTGGCAGCAGGATC-3'<br>5'-CCAATCCTTATTAAGGCTGGCGATATTATTTATC-3'<br>5'-AGCTCGATATCGCATGATTTTGTTCTGTTCAAACAGATTATTC-3' | This study |
| Deletion of *ttdU* in *S*. Tm | 5'-CTAGAGGTACCGCATGCTCGCGGATAGCCCAGGT-3'<br>5'-ATGGGATATGGGAGAAATTATTCCCATACAAAAACTAAGTC-3'<br>5'-TTTCTCCCATATCCCATATGACCGACGTTATG-3'<br>5'-AGCTCGATATCGCATGAAGCGGGTCACACCTTCA-3' | This study |
| Deletion of *oadGAB* in *S*. Tm | 5'-CTAGAGGTACCGCATGCGTGACCATCGCGTTGCA-3'<br>5'-GGCTATTTTTCCAATCTAATCGGGAGCAGT-3'<br>5'-ATTAGATTGGAAAAATAGCCTTCAGGCA-3'<br>5'-AGCTCGATATCGCATGAAAGAGTTTATTGCGCAGA-3' | This study |
| Deletion of *frdABCD* in *S*. Tm | 5'-CTAGAGGTACCGCATGGCGTTGGCGTGCGAGAGA-3'<br>5'-ACCCAGAGCCTGCCTCTCATTTTCCCGC-3'<br>5'-GAGGCAGGCTCTGGGTTCTGCGAAGTATG-3'<br>5'-AGCTCGATATCGCATGACTGGTCATTAAATAGAGGTTGATC-3' | This study |
| Deletion of *ttdABT* in *E. coli* | 5'-GCTTCTTCTAGAGGTACCGCATGCAAAACTGCAACCAATGC-3'<br>5'-CCGTCCTCACTTTGTCTGGAAAAATTCACC-3'<br>5'-TCCAGACAAAGTGAGGACGGTGCCGGAA-3'<br>5'-GGAGAGCTCGATATCGCATGTGGCGGCTCAGGGTACTG-3' | This study |
| Complementation of *ttdBAU* in *S*. Tm | 5'-GCTTCTTCTAGAGGTACCGCATGTTATTTGATGAAACTTACGTTCTGGC-3'<br>5'-GGAGAGCTCGATATCGCATGCTTTTCGCCTGCGTGAGC-3' | This study |
| Complementation of *dtuT* in *S*. Tm | 5'-CTTCTTCTAGAGGTACCGCATGCAGACAGATTTTAAGAGGAAATTATTTG-3'<br>5'-GGAGAGCTCGATATCGCATGTTAATAGAAGGGATAAAATATCGG-3' | This study |
